# Microfluidics for Electrophysiology, Imaging, and Behavioral Analysis of *Hydra*

**DOI:** 10.1101/257691

**Authors:** Krishna N. Badhiwala, Daniel L. Gonzales, Daniel G. Vercosa, Benjamin W. Avants, Jacob T. Robinson

## Abstract

The cnidarian *Hydra vulgaris* provides an exciting opportunity to discover the relationship between animal behavior and the activity of every neuron in highly plastic, diffuse network of spiking cells. However, *Hydra’s* deformable and contractile body makes it difficult to manipulate the local environment while recording neural activity. Here, we present a suite of microfluidic technologies capable of simultaneous electrical, chemical, and optical interrogation of these soft, deformable organisms. Specifically, we demonstrate devices that can immobilize *Hydra* for hours-long simultaneous electrical and optical recording, and chemical stimulation of behaviors revealing neural activity during muscle contraction. We further demonstrate quantitative locomotive and behavioral tracking made possible by confining the animal to quasi-two-dimensional micro-arenas. Together, these proof-of-concept devices show that microfluidics provide a platform for scalable, quantitative cnidarian neurobiology. The experiments enabled by this technology may help reveal how highly plastic networks of neurons provide robust control of animal behavior.

## INTRODUCTION

Understanding the relationship between animal behavior and the activity of individual cells in the nervous system would be a major scientific breakthrough. To reach this goal, scientists are developing new electrical and optical technologies capable of simultaneously recordings from hundreds of individual neurons with the temporal resolution to capture individual action potentials.^1–15^ These technologies, however, fall well short of recording every action potential from each individual neuron in vertebrate model organisms that have neurons numbering from the hundreds of thousands to tens of billions.

Thus, to observe cellular level activity of the entire nervous system, scientists turn to small invertebrates like *Caenorhabditis elegans* and *Drosophila melanogaster*. In addition to having far fewer neurons, their small size and transparency facilitates in vivo calcium- or voltage-sensitive fluorescence imaging that can record simultaneous activity of hundreds to thousands of individual neurons.^11,16–19^ To make these investigations even more attractive, several lab-on-a-chip technologies now provide increased throughput for chemical, optical, and electrical interrogation of *C. elegans* and Drosophila on microfluidic platforms. This confluence of technologies has revealed how many behaviors can be implemented by neural circuits,^20–24^ however, *C. elegans* and *D. melanogaster* may not be the best suited to study neural circuit repair and remodeling. Although neurites connecting cells can regrow when severed, if even a single neuron is ablated, *C. elegans* or *D. melanogaster* often show significant and permanent behavioral deficits.^21,23,25–31^ This static and fragile neural architecture stands in stark contrast to the mammalian cortex, which can remodel itself to retain or regain function despite the loss of a significant number of neurons.^32–34^

In contrast to *C. elegans* and *D. melanogaster*, the architecture of the *Hydra* nervous system is extremely dynamic making it an exciting model for studying neural plasticity and repair. While the *Hydra* are small (0.5 - 15mm in length) and transparent like *C. elegans* and *D. melanogaster* larvae, the entire population of neurons in *Hydra* nervous system is continually replenished and the number of neurons can vary by more than a factor of ten depending on nutrient availability. Despite these dramatic changes in the number of neurons and the lack of static structures, the animal maintains stereotypical behaviors.^35–37^ Moreover, *Hydra* can rapidly repair itself following a sudden loss of neurons. When the animal is bisected, the organism reforms and resumes natural contractile behaviors in as little as 48 hours due to high differentiation capability of the stem cells.^36,38,39^

*Hydra* are also a compelling model organism because their diffuse network of spiking neurons resembles neural network models often studied by computational neuroscientists.^40^ *Hydra* have several genes that encode voltage-gated ion channels allowing their neurons to generate fast action potentials similar to those in mammalian nervous systems.^41^ Ultrastructural studies implicate both electrical and chemical synapses in *Hydra* along with some common neuropeptides and neurotransmitters.^42–48^ Thus, unlike *C. elegans*, whose neurons lack sodium driven action potentials, *Hydra* (like *D. melanogaster*) have genes encoding for voltage-gated sodium channels and thus provides opportunities to study information processing in simple networks of spiking neurons.

While the small size of *Hydra* offers several advantages as a model organism, it also presents challenges for moving and manipulating the organism and delivering well-controlled stimuli. In the case of *Drosophila* and *C. elegans* that are similarly sized, this challenge has been addressed using microfluidic technology.^49–51^ Microfluidics provides robust and scalable methods to reversibly restrain and physically manipulate *Drosophila*^52–56^ and *C. elegans.*^57–67^ Specifically, in the case of *C. elegans*, microfluidics have been shown to provide precise control over the local environment for observing taxis and locomotive behaviors, performing calcium imaging and recording electrophysiological activity from the pharynx and body-wall muscles.^22,61,65,68–71^

Unfortunately, direct application of the existing microfluidic technologies is unlikely to be successful with *Hydra* due to its soft and deformable body. Unlike *C. elegans* and *Drosophila, Hydra* has neither a tough protective cuticle nor a stereotyped size. Miniscule forces, on the order of nano-newtons, are sufficient to tear the epithelial cell layers to form an oral cavity. Body contractions themselves can generate forces of this magnitude.^72^ Thus, the spontaneous body contractions and elongations can shear and dissociate the epithelia, if the aggressive microfluidic confinement strategies successful in small invertebrates like *C. elegans* are translated directly to *Hydra*. Furthermore, within a clonal population, *Hydra* may vary in size by more than a factor of ten and an individual animal can change length by an order of magnitude during contraction. Thus, any microfluidic confinement or immobilization strategy must accommodate deformable animals of a variety of sizes and reduce shear forces.

Here we show that specially designed microfluidic devices enable key neurobiological experiments to be performed in *Hydra*. Specifically, we illustrate safe handling and manipulation of the gelatinous *Hydra* in a microfluidic environment for several hours to days by carefully controlling fluidic pressure. We also show how the microfluidic devices allow us to use electrical and optical techniques to simultaneously measure the of activity of muscle cells and the group of neurons responsible for motor function during body column longitudinal contractions. We can also stimulate specific behaviors, such as feeding, by using chemical stimulants to study the cellular level activity at the onset and during the behavior. We also replicate and quantitatively analyze a subset of the *Hydra* behaviors in the microfluidic arena essential for behavioral and locomotion assays. This is believed to be the first microfluidic platform for manipulating *Hydra* for scalable behavioral and neuroscience studies.

## RESULTS AND DISCUSSION

### Manipulation and immobilization of *Hydra* in microfluidic devices

Despite the soft and deformable body of the *Hydra*, we found that with care, we could transfer the animal into and out of the microfluidic devices with roughly 95% success (Fig. 1a,b, Supplementary Movie 1). To load *Hydra* into a microfluidic device, we use a transfer pipette to move an animal into an open syringe cap connected to the device via tygon tubing. Because *Hydra* readily attach themselves to many surfaces, we found that drawing the animal into the tygon tubing quickly dramatically improved the success of loading. Once loaded, the animal’s position can be precisely controlled with gentle application of fluid flow. Unloading follows a similar procedure in reverse (see Methods).

**Figure 1:**
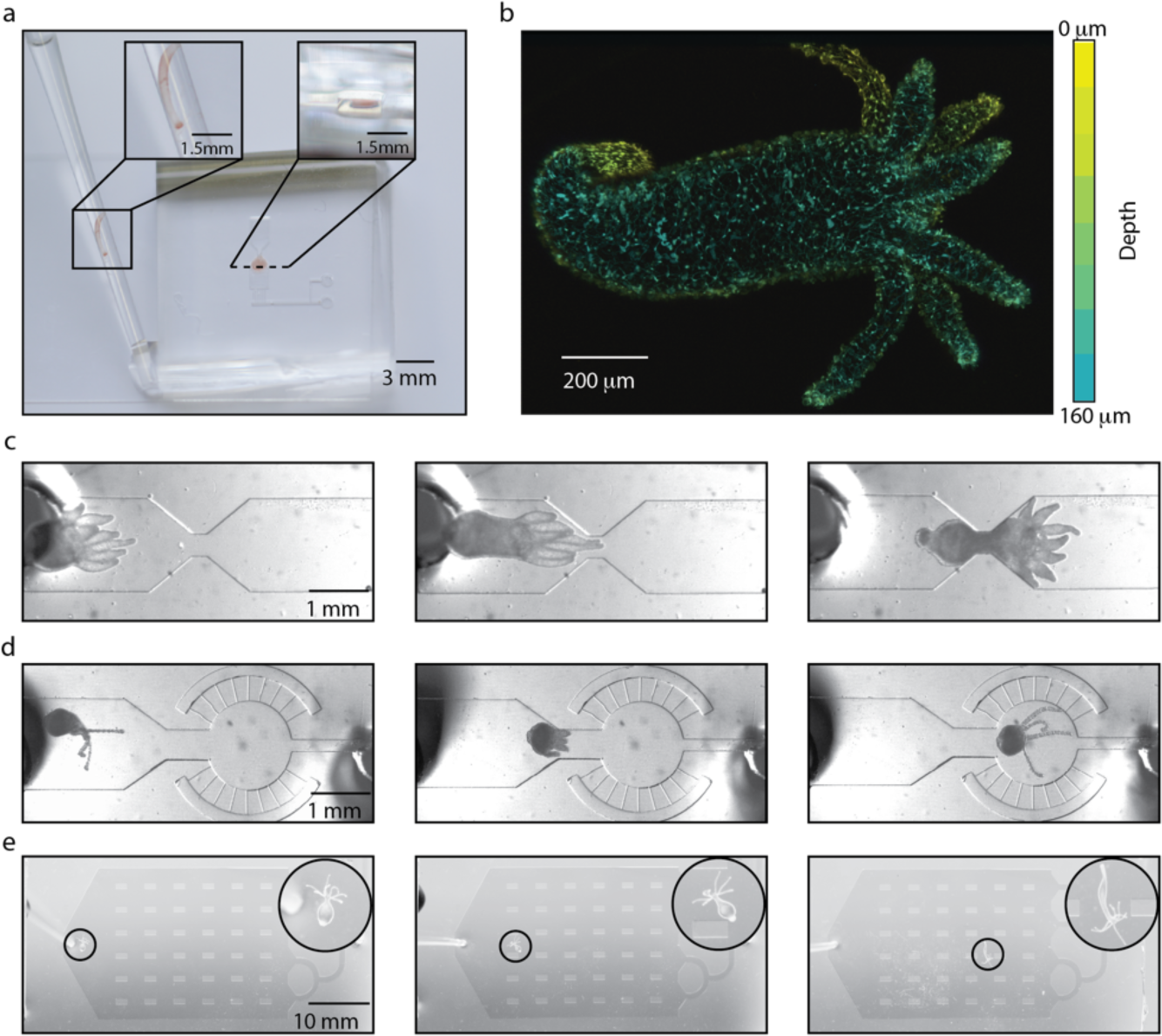
Microfluidic Manipulation and Immobilization of *Hydra*. **(a)** Typical *Hydra*, 0.5 mm in diameter, is immobilized in a 110 μm tall hour-glass microfluidic chamber. For *Hydra* immobilization and size comparison, inset on left shows zoom-in of freely moving *Hydra* in a 1 mm diameter glass pipette. Inset on the right shows zoom-in of a cross-section of microfluidic chamber with similar sized flattened *Hydra*. **(b)** Maximum fluorescence intensity projection image shows *Hydra* nerve net with the colors (yellow to teal) corresponding to depth (from z-stack taken with confocal microscopy). *Hydra* with pan-neuronal expression of GFP under actin promoter was immobilized in a 160 μm tall microfluidic chemical interrogation chamber and anesthetized with 0.1% chloretone on-chip prior to imaging. **(c-e)** Optical micrographs show loading (left), recovery (middle) and precise positioning of *Hydra* (right) in three different microfluidic chambers: **(c)** hour-glass chamber for electrophysiology constraining the body column from large movements; **(d)** wheel-and-spoke perfusion chamber constraining locomotion; **(e)** behavioral micro-arena with 2 x 1 mm micro-pillars limiting movements and locomotion in the vertical plane but allowing all movements in the horizontal plane. Inset: zoom-in of behaving *Hydra* in the micro-arena.

To showcase how microfluidics enable a variety of *Hydra* studies ranging from electrophysiology to quantitative analysis of locomotion, we created three classes of immobilization chambers: 1) hour-glass shaped chambers that reduce *Hydra* movement for high-resolution cellular imaging and electrophysiology (Fig. 1c); 2) wheel-and-spoke geometries that confine *Hydra* to a region roughly the size of the animal to facilitate imaging and chemical perfusion (Fig. 1d); 3) open-field geometries that allow *Hydra* to move and explore a quasi-2D environments (Fig. 1e). For each immobilization chamber, we performed proof-of-principle experiments to demonstrate how these devices can help study *Hydra* neural activity and/or behavior.

**Figure 2:**
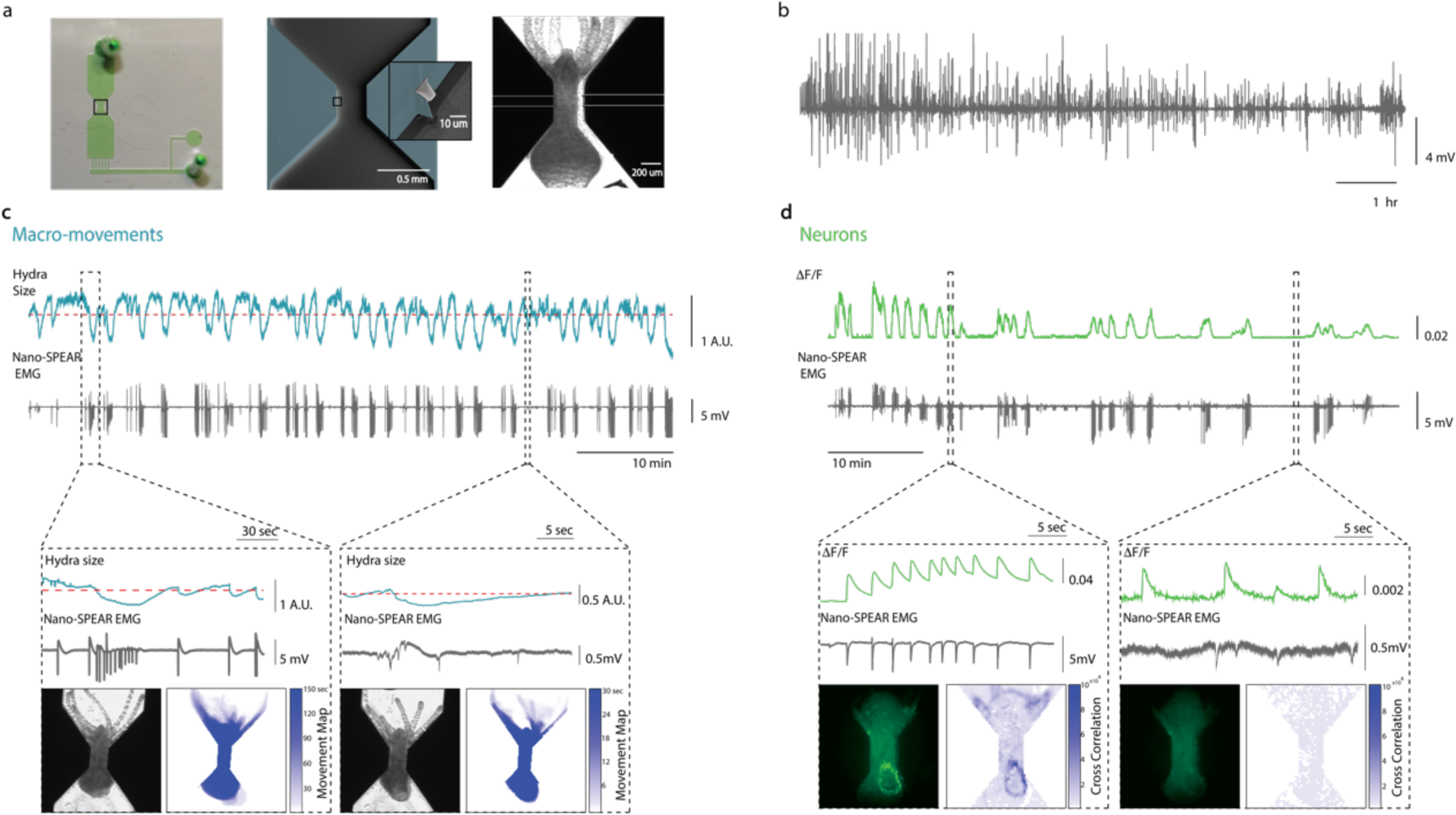
Simultaneous electrophysiology and calcium imaging. **(a)** Photograph (left) of a microfluidic immobilization chamber filled with green dye. Black box highlights the recording region in the microfluidic chamber (110 μm tall). False colored scanning electron micrograph (middle) shows the recording region (blue, photoresist; light grey, Pt; dark grey, silica) on the nano-SPEAR chip (50 μm tall). Inset shows a zoom-in of the Pt electrode (light grey) suspended mid-way between the top and bottom of the photoresist sidewall(blue). Optical micrograph (right) shows *Hydra* immobilized in the microfluidic chamber placed on top of the nano-SPEAR chip with combined 160 μm tall recording region. **(b)** Representative 10-hour– ong electrophysiology trace measured from *Hydra* vulgaris AEP. **(c, d)** Simultaneous electrophysiology and imaging from **(c)** *Hydra* (H. Vulgaris AEP) and **(d)** neurons in transgenic *Hydra* (GCaMP6s, neurons) (n=1). Top trace shows c) change in body size (area) of the top half of the *Hydra* body (10 Hz) and (d) mean fluorescence (ΔF/F) neurons (20Hz). Bottom trace shows simultaneously recorded electrical activity from the muscles using nano-SPEARs electrode (1KHz). Left box show correlation during high activity period, contraction burst (c, d). Right box shows correlation during low activity period, (c) tentacle activity or (d) RP-like activity. Within each box, top traces show (c) change in body size, (minima in body size trace means contractions) or (d) peaks in fluorescence (during contractions) coinciding with peaks in electrophysiology (bottom trace). (c) A representative micrograph (left) of *Hydra* shows representative body size. The movement map (left) shows the body regions moving during the period. (d) A representative fluorescence micrograph (left) shows calcium levels with high fluorescence. Correlation map (right) spatially plotting the correlation coefficients shows the activity from region in *Hydra* body with high correlation with electrophysiology.

**Figure 3:**
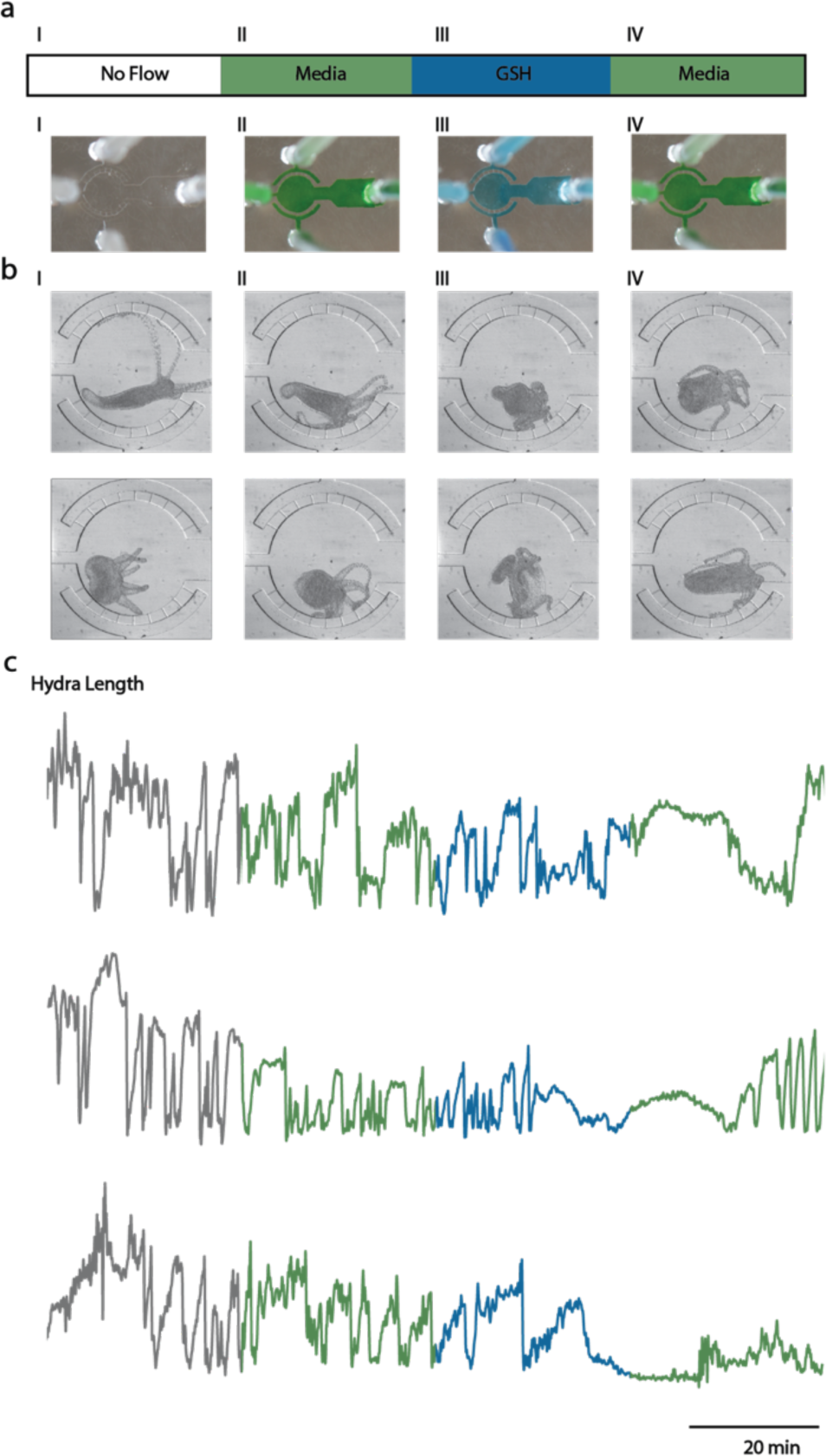
Functional imaging during chemically induced feeding behavior with reduced glutathione (GSH). **(a)** Photographs show a microfluidic chamber with a circular observation chamber in the middle surrounded by small perfusion channels. The narrow perfusion channels with a port on the top and bottom are used for perfusion inflow and outflow, respectively. The dye colors indicate flow conditions: (I) no flow (clear or grey); (II) flow of *Hydra* media (green); (III) 9µM GSH (blue); (IV) *Hydra* media (green). **(b)** micrographs of immobilized *Hydra* show representative activity: (I, II) stereotypical body elongations (top) and contractions (bottom) with no perfusion flow or with media flow; (III) feeding response induced by perfusion of GSH leading to tentacle writhing (top) and outward folding of the epithelia from mouth opening (bottom); (IV) Recovery of typical body contractions (top) and elongations (bottom) following perfusion of *Hydra* media. **(c)** Body length traces show contractions and elongations from three individual *Hydra* (colors correspond to flow conditions I-IV). Minimum body length indicates contraction.

### Electrophysiology and imaging of immobilized *Hydra*

The hour-glass tight confinement chambers immobilize the animal against the walls of the microfluidic device allowing us to minimize *Hydra* movement for *both* high-resolution optical imaging *and* cellular-scale electrophysiology using nano-SPEAR electrodes that protrude from the walls of the microfluidic channels.^71^ These hour-glass chambers effectively immobilize the animal by first flattening the deformable *Hydra* in the roughly 110 µm tall microfluidic channels and then pinch a portion of the mid-body column of the animal to keep it immobile (Fig. 1c). We found that these chambers reduced the movement of *Hydra* cells in the entire body column to approximately 65 um/minute (see Methods), though the tentacles were largely free to move. A major advantage of these hour-glass shaped immobilization chambers (based on previously reported immobilization chambers for *C. elegans*)^58,71^ is that they avoid sharp corners that can damage the soft *Hydra* body and they can accommodate the large differences in animal sizes found in the *Hydra* colonies (Fig. 1c).

Using our unique ability to perform simultaneous electrophysiology and imaging in intact *Hydra*, we sought to identify the origin of the electrical signals recorded from the *Hydra* body. The *Hydra* body is mostly comprised of two layers of contractile epitheliomuscular cells (20-40µm in length) capable of generating action potentials and innervated by a smaller number of neurons (8-10µm) that also are believed capable of generating action potentials.^73^ Using this tight confinement device, we were able to record electrical activity from a single animal for 10 hours using nano-SPEAR electrodes. In these recordings, we observed a mixture of high and low amplitude electrical spikes (Fig. 2b). We then performed simultaneous brightfield imaging in *Hydra vulgaris* AEP for 1 hour under dark conditions, and observed strong correlation between body and/or tentacle contractions and the electrical spikes recorded with our nano-SPEARs (Fig. 2c). This correlation suggests that the electrical signals primarily represent action potentials generated by the muscle cells, which is consistent with previous recordings using nanoSPEAR electrodes in *C. elegans.*^71^ To quantify the body and tentacle contractions we measured the area occupied by the *Hydra* in the upper and lower halves of the microscope image (body size). We found that the majority of small amplitude electrical spikes correlated with tentacle contractions (or small amplitude changes in size of the upper body half). Small amplitude spikes were also observed in the absence of obvious tentacle contractions suggesting that nanoSPEARs may also record spiking activity from other cells. The large amplitude waveforms coincided with body contractions bursts (CB) (or large amplitude changes in the size of both the top and bottom halves of the animal) (Fig. 2c) (Supplementary Movie 2). This pattern of small and large amplitude waveforms was observed across six individual animals (Supplementary Fig. 1). Together, the absence of high amplitude electrical activity during body-elongations, when the rhythmic potential (RP) neurons are thought to be active,^35^ and large percentage of electrical activity measured during body or tentacle contractions further indicates nano-SPEAR electrodes predominantly measure ectodermal muscle activity associated with body or tentacle contractions.

Having determined that the electrical signals recorded from our nano-SPEAR electrodes represent muscle activity, we then looked for the neural activity patterns that drive muscle contractions. By performing simultaneous electrophysiology of the muscle cells and calcium imaging in neurons (using a transgenic strain that expresses GCaMP6s pan-neuronally), we could correlate neuronal activity with muscle contractions. When we compared this simultaneously recorded muscle and neuronal activity, we found that body column contractions were driven by a nerve ring in the *Hydra* foot, and that tentacle contractions were modulated by neurons in each tentacle (consistent with previous reports^35^) (Fig. 2d). Specifically, during body column contractions calcium imaging showed synchronous firing of the cluster of neurons in the nerve ring at the foot. Synchronized with this calcium activity, we recorded large amplitude electrical spikes from the epithelial muscle cells (Supplementary Movie 3). Computing the cross-correlation between calcium-sensitive fluorescence imaging and electrophysiology during contractions shows that the neurons in the foot indeed correspond to body contractions. All tentacles also contract during contraction bursts and we also see strong correlation of electrical activity with the activity of neurons located near the base of the tentacles. Interestingly, when the body column is elongating or stationary we find little correlation between neural activity and electrically-detected muscle activity. Thus, the RP neurons that are active during elongation appear unassociated with any muscle activity (Fig. 2d, right) suggesting that they may play a role in inhibiting body column contraction. During these periods, we often measure isolated, very low amplitude spikes in the electrical activity though neither the pattern nor the timing was correlated well with RP neuronal activity (Fig. 2d, right).

We found that for the hour-glass immobilization chamber, experiments could last between 1 and 10 hours depending on the experimental conditions allowing us to observe many cycles of contraction and elongation. The excitation light used for fluorescence imaging stimulated inchworm-like locomotion away from the recording site and as a result typical imaging experiments could last roughly one hour. Under low-light conditions during electrophysiology experiments, the animal was much less motile, and experiments could typically last more than 10 hours.

### Chemical stimulation of *Hydra* in microfluidics

Chemical stimulation is key tool for neurobiologists allowing them to trigger precisely timed behaviors or to investigate the role of neuromodulators and/or ion channels using known agonists or blockers. To apply chemical stimulation to *Hydra* without stimulating a mechanical response to changing fluid flow rates, we developed 160 μm tall wheel-and-spoke perfusion chambers (Fig. 1d, 3). A key design element of these chambers is a slow perfusion rate that avoids stimulating natural responses to changing fluidic pressures or sheer stress. We found that high flow velocities in large microfluidic channels often initiated body contractions or tentacle swaying. At times, high flow rates produced sheer forces that damaged *Hydra*. We also observed that *Hydra* would frequently bend or translocate in the direction of the flow. Thus, to apply chemical stimuli without initiating these behaviors, we created microfluidic devices that minimize the flow rates of chemical stimuli. To minimize the rate of fluid flow into the *Hydra* observation chamber, we relied on the large fluidic resistance created by short and narrow (25 x 20 um) perfusion channels leading into the larger observation chamber (1500 radius × 160 height, μm). This geometry is based on the previously reported chemical perfusion chambers for perfusion in cell culture arrays.^74^

Proof-of-concept experiments show that perfusion through our wheel-and-spoke chambers induce chemically stimulated behaviors without evoking mechanical responses to fluid flow. During 30-minute periods of no fluid flow (I), *Hydra* media flow (II), reduced glutathione (GSH) flow(III), and recovery with *Hydra* media flow (IV), we observed feeding responses as expected with exposure to GSH but no change in behavior during *Hydra* media flow. When we perfused *Hydra* media through our chemical stimulation chamber, we observed minimal change in contraction rate or bending suggesting that the flow rates were sufficiently low to prevent mechanical stimulation (Fig. 3a-c II). Contractions and elongations were quantified by measuring the decreases and increases in body length, respectively (see methods). We found no obvious difference in the contraction rates either with or without the flow of *Hydra* media (Fig. 3b,c I-II, Supplementary Movie 4). The continuation of this periodic body contractions and elongations pattern during flow indicated the perfusion flow rates caused negligible mechanical stimulation.

Because the low fluid flow rates engineered in our wheel-and-spokes chambers do not significantly affect *Hydra* behavior, we were able to perfuse GSH (9uM) to induce a feeding response and inhibit body contractions (Fig. 3a-c III). Normal body contractions were interrupted after approximately 15-20 minutes of GSH flow as seen by the lack of sharp decreases in body length as expected through previous reports^75^ (Fig. 3c III) and followed by tentacle writhing (Fig. 3b III, top). During tentacle writhing, body length remained constant but tentacles contracted and curled towards the mouth until the oral cavity was formed (Fig. 3b III, bottom). Once the oral cavity had formed, epithelium began folding outwards increasing the mouth size until *Hydra* lost its tubular shape and hydrostatic rigidity (Supplementary Movie 4). The chemically induced response in *Hydra* was then reversed by switching the perfusion input from GSH to *Hydra* media. In all three trials, we successfully recovered the contractile activity as seen by the return of spikes in the body length of *Hydra* (Fig. 3c IV). In all trials, except one, the folded *Hydra* body eventually regained its tubular shape by resealing the oral cavity through the duration of the trial.

The ability to chemically stimulate *Hydra* feeding responses in microfluidic chambers provides the exciting opportunity to image neural activity during these behavioral state transitions. As an example, we imaged neuronal activity in transgenic *Hydra* (GCaMP6s, neurons) under GSH stimulation of the feeding response. To reduce the effects of photobleaching we imaged *Hydra* for five minutes under the flow of *Hydra* media, followed by thirty minutes of GSH, and a second thirty-minute period of *Hydra* media flow (Supplemental Movie 5). We observed that during GSH stimulation, normal body contractions were interrupted and the cluster of neurons in the foot became less active after approximately twenty minutes of GSH flow (Supplementary Movie 5). The neurons at the base of tentacles were active during tentacle writing and the neurons in the upper half of the body seemed more active during mouth opening. During the recovery period with *Hydra* media flow, we initially observed a smaller subset of neurons in the lower half of the body become active but did not lead to body contractions until larger ensemble of neurons in the foot, likely belonging to contraction burst circuit, became active. This ability to perform cellular-resolution imaging during chemically stimulated behaviors provides new opportunities to study sensory-motor transformations in the entire network of spiking neurons in *Hydra.*

### Behavioral Analysis of *Hydra* in microfluidics

The quasi-2D environment provided by microfluidics also helps us quantitatively track *Hydra* locomotion and body posture as they explore their surroundings. In addition to periodic body contractions and elongations, *Hydra* can also explore their environment by bending and swaying or move to new locations through inch-worm, somersault or swimming locomotion. Because microfluidic arenas reduce *Hydra* movements to a quasi-2D plane, the task of quantifying *Hydra* movements and posture is greatly simplified and we can use a simple, low-cost camera placed above the device. Microfluidics also provides an excellent platform for controlling chemical, thermal, and physical conditions. The combination of behavioral tracking and well-regulated environmental conditions will help reveal sensory-motor processing in these simple neural networks.

We found that in microfluidic devices roughly 3 cm wide with channel height’s ranging from 100 – 500 µm *Hydra* display similar behaviors to unrestrained animals in 3D including contraction, elongation, swaying, as well as inch-worm and floating locomotion. Based on our observations of *Hydra* over several hours in our 440µm tall behavioral arenas (Fig. 1e), we classified the *Hydra* behavior into two broad behavioral classes: exploration and locomotion. We found that just like in flask cultures, *Hydra* in our behavioral arenas typically anchors itself to the top or bottom surface with its foot while periodically contracting and elongating (Supplementary Movie 6). Less frequently, we observed swaying or bending, which is also seen in flask cultures. In addition to these exploratory behaviors we also observed *Hydra* locomotion by either inchworm or floating. We did not observe somersaulting as has been reported in 3D environments^76^, perhaps due to the short height of the quasi-2D chamber or the relative rarity of somersaulting events.

By reducing body posture and locomotion to 5 variables, we could quantify *Hydra* behavior over several days (Fig. 4). We defined the change in body length of *Hydra*, L, which allows us to easily detect body contraction events (Fig.4 a, Supplementary Fig. 2). Because of the low imaging frame rates used, we identified contraction burst (CB) events rather than individual contraction burst pulses (Fig. 4c). On average, we measured similar rate of 9 -15 CBs per hour as previously reported.^77^ We defined body orientation as a, the angle of a vector from the foot to the mouth with respect to the positive x-axis. However, we noted that body column of *Hydra* often curved. The body orientation, α, did not represent the direction of the mouth especially when *Hydra* body formed a U-loop. Thus, we obtained body posture by fitting two vectors separated along the midpoint in the body column from the foot to the center of the body and from center of the body to the mouth, α_1_ and α_2_. The difference between these two angles, b, provided information about the curvature along the body column. In cases when the body column was straight, β was nearly zero. When the body column had slight curvature, there was a small difference between α_1_ and α_2_, β < π/3. Similarly, when *Hydra* body looped to form U-shape, the angular difference between the two vectors was large, π > β >= π/3 (Fig. 4a, c). Alternatively, comparing the difference in body length based on the Euclidean distance between the mouth and foot either with or without accounting for the body center further confirmed the bending events. Thus, by segmenting the *Hydra* body, we were able to gain additional information about the body postures. Interestingly, the U-shaped posture frequently occurred following translocation events.

**Figure 4:**
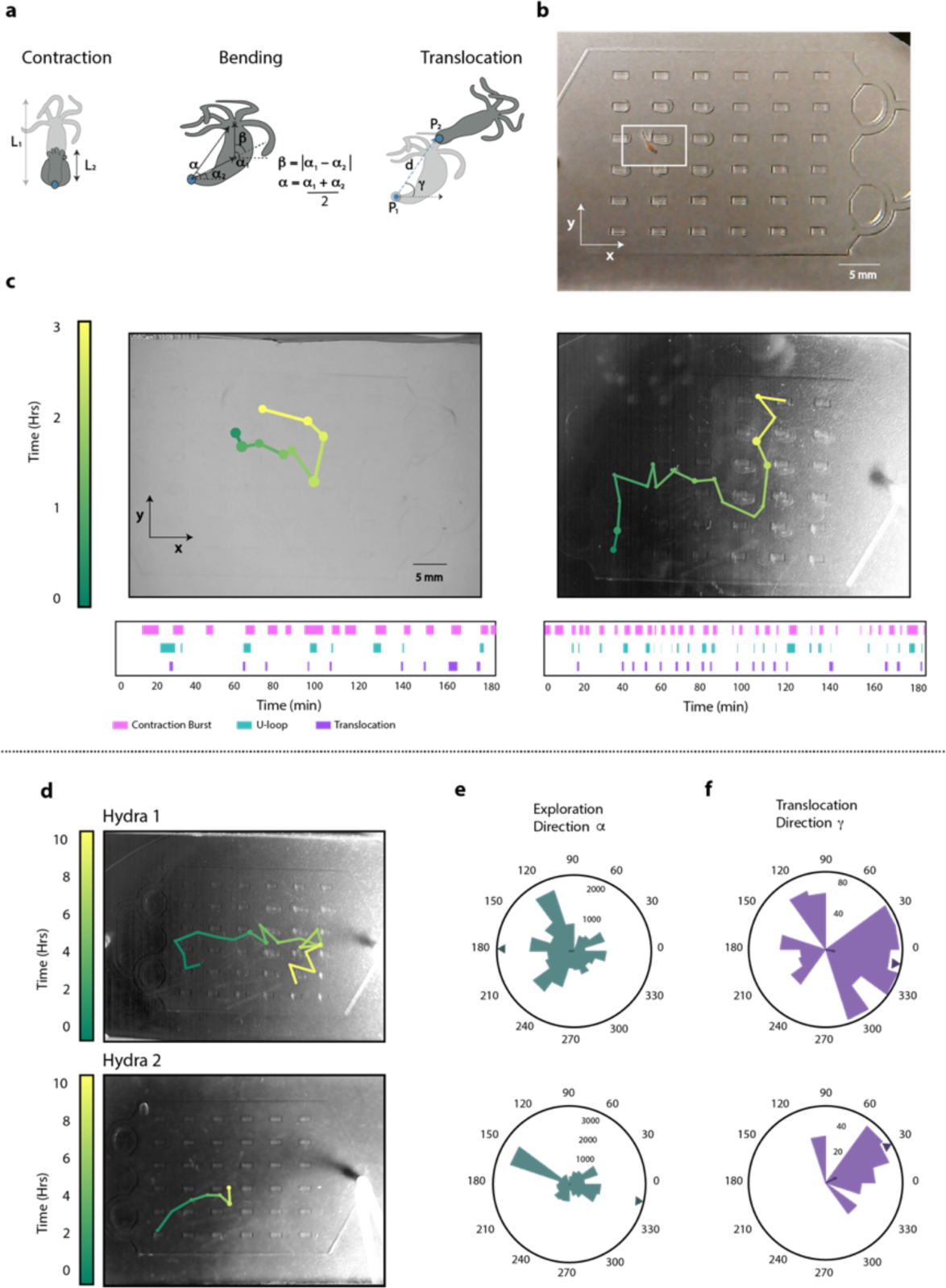
Behavioral Analysis in Microfluidic Arenas. **(a)** Schematics show common behaviors of *Hydra* observed in microfluidic arenas: body contractions (periodic decrease in length of the body, L), exploration (reorientation of the head, α), bending (body curvature and turning head towards the foot, β), and translocation (distance and direction of displacement of the foot to new location, d and γ). **(b)** Photograph shows microfluidic arena for locomotion and behavioral observation of immobilized *Hydra* positioned with uniform lighting (top). **(c)** Skeletonized track (3 hours) of *Hydra* movement manually tracked in a microfluidic arena with no light gradient (left) and with light gradient (right). The size of each node corresponds to the time spent at that location. The length of each edge corresponds to the distance traveled during a translocation event. The raster plots show when each of the tracked behavior is occurring during the recording period. **(d-f)** Phototactic behavior in microfluidic arenas with a light gradient. **(d)** Skeletonized tracks (10 hours) of two different *Hydra* under a light gradient. **(e)** Distribution of body orientation (exploration direction) in a light gradient during the 10-hour imaging. **(f)** Distribution and preference for direction of the *Hydra* translocation in a light gradient. Bold arrows show the magnitude of resultant vectors and the triangles near the outer circle clearly indicates the preferred direction for **(e)** exploration and **(f)** translocation.

Using time-averaged location of the foot for generating the movement track of *Hydra* through the microfluidic arena, we found the displacement events were significantly less frequent compared to the body contractions (Fig. 4c). *Hydra* periodically contracts and elongates in a variety directions while the foot is anchored to a single location. Translocation occurs when the *Hydra* releases the foot and reattaches it at a different location. Thus, tracking the location of the foot shows the locomotion pattern (Fig. 4c) of *Hydra* where the distance, d, and the direction traveled, γ, are represented by the lines on the track. The locomotion steps were frequently stereotyped inchworm movements where *Hydra* expanded its tentacles and contracted the rest of the body towards the tentacles. *Hydra* was also seen bending over its tentacles to complete the locomotion step. Often *Hydra* took several steps before reattaching the foot. For simplicity, we identified the position where the *Hydra* foot finally readhered as the step final position to determine the locomotion step size and direction (Fig. 4c). Under uniform light condition (n=1), locomotion of *Hydra* seemed random with no clear preference for direction (Fig. 4c left).

We can also use these microfluidic arenas to study more complex locomotion patterns that are influenced by external factors such as light. By observing *Hydra* locomotion for several hours in the microfluidic devices, we found *Hydra* movement in a light gradient resembles a directed random walk toward the brighter end of the device (Fig. 4c (right), d). We observed this behavior both when the microfluidic chip was positioned vertically and horizontally. We quantified the directed random walk using 5 different *Hydra* cultured in dark environments. For each experiment, we transferred *Hydra* into the microfluidic chip at least 24 hours after feeding (n=5). When we averaged all the translocation events for two representative *Hydra*, we found a resultant vector in the direction of the bright side of the chip (Fig. 4f). While this data is consistent with previous reports of positive phototaxis in *Hydra*,^78^ more work is needed to better understand this behavior. For example, we observed clear preference towards the light during the first 24 hours in the device in *Hydra* that showed multiple translocation events; however, some animals remained in the same place for the majority of the recording period or after days of immobilization in the arena exhibited negative phototaxis. We also found *Hydra* did not always have preference for orienting its body towards the bright side of the chip (Fig. 4e). This finding indicates that there may be several elements influencing phototaxis in *Hydra* and that these behaviors may be influenced by circadian factors. The fact that we can quantify behaviors such as locomotion and we can alter the environmental conditions in these microfluidic arenas strongly suggests we can study the underlying mechanisms for sensory motivated behaviors, such as phototaxis, in *Hydra*.

## DISCUSSION

Just as microfluidic technologies have accelerated studies of *C. elegans* and *Drosophila*,^22,49,51–54,58–62,71,79–81^ we believe that the electrophysiology, imaging, environmental control, and behavioral tracking made possible with the microfluidic technologies shown here will help *Hydra* become a more powerful model organism. In particular, we have found that despite the soft and deformable body of *Hydra*, we can create microfluidic chambers than can immobilize the animal with varying levels of confinement. The easy loading and unloading protocol means the same *Hydra* can be studied over several days. Moreover, the microfluidic platform allows gentle perfusion of chemicals and buffers for stimulating behaviors or studying locomotive behaviors in complex chemical, thermal, or optical landscapes.

As a demonstration of the types of experiments enabled by microfluidic immobilization, we showed simultaneous electrical recordings from muscles and optical recordings from the neurons, providing insight into the patterns of neural activity that drives body column and tentacle contraction. We see this type of combined electrophysiology and whole-brain imaging as a powerful method to study coordination between neural activity and body movements - a key step toward decoding neural activity.

The *Hydra* microfluidic platform enables chemical stimulation of behavior similar to those used to probe neural circuits in other invertebrates.^69^ Although *Hydra* are highly sensitive to fluid flow, we could reduce the perfusion to sufficiently slow rates and minimize the responses of *Hydra* to this mechanical stimulus. These types of experiments may help us understand the neural circuits that process external stimuli and execute resulting motor programs.

In addition, the quasi-2D environment provided by microfluidics makes it easier to quantify the *Hydra* posture and movements and facilitates whole-brain calcium imaging. Axial scanning during optical microscopy is typically the slowest scan axis because it typically requires moving an objective lens or sample stage. Thus, by confining *Hydra* to a plane less than 200 µm thick will increase the rate at which one can acquire whole-brain imaging data.

Overall, the seemingly simple cnidarian *Hydra* combined with a microfluidic interrogation platform provides many opportunities to discover how complex behaviors are implemented in dynamic networks of spiking neurons. For example, the microfluidic environments described here could be used to combine whole-brain imaging with locomotion and directed movements like chemo-, or photo-taxis. Integrating heating elements^82^ or microactuators,^83^ could extend these investigations to cover thermo- or mechano-sensory processing. Because *Hydra* survive for days in these microfluidic chips, we envision that these behavioral screens could be performed with many animals in parallel over extended periods of time.

Through studies like these enabled by our microfluidic platform, it may be possible to understand simple rules governing the function of highly plastic neural circuits that may be conserved in more complex brain architectures.

## MATERIALS AND METHODS

### Device Fabrication

The microfluidic chips were fabricated using approximately 5 mm thick layer of polydimethylsiloxane (PDMS) (Sylgard) molded from SU-8 2075 (MicroChem) master mold. The microfluidic chip for electrophysiology was molded from ~110 µm thick features on the master. The chemical perfusion chip was molded from a master mold fabricated with two layers, ~25 µm perfusion channels with SU-8 3025 and ~150 µm observation chamber with SU-8 2075. The behavioral microfluidic arena was fabricated from ~440 µm tall features containing 2 x 1 mm pillars on the master. The ports for inserting tubing were punched with either 1 or 1.5 mm biopsy punches. Larger biopsy punch was used for *Hydra* insertion port. The microfluidic chips for chemical perfusion and behavioral arena were permanently bonded to glass wafer with oxygen plasma treatment at 330 mTorr for 30 sec. The microfluidic chips for electrophysiology and imaging were clamped to the nano-SPEAR chip with a customized acrylic enclosure. Clamping with acrylic instead of permanently bonding PDMS to the electrophysiology chip makes cleaning the device easier between uses.

The electrical interrogation device is a combination of electrophysiology chip interfaced with a microfluidic chip similarly to the previously reported nanoSPEARs chip.^71^ The electrophysiology device was fabricated on a glass substrate (University Wafers) using micro- and nano-fabrication techniques. A layer of KMPR photoresist was spun on the glass wafer. Platinum electrodes were patterned on the first KMPR layer. A second layer of KMPR was patterned with the immobilization chamber on top of the Pt electrodes using photolithography. The recording chambers patterned on the top KMPR layer were then etched to the bottom KMPR layer with reactive ion etcher (RIE) resulting in suspended electrodes. To increase the longevity of the device, the entire device was coated with Parylene C which acts as a water barrier. The tips of the electrodes were exposed with focused ion beam milling. KMPR has high autofluorescence which contributes to high background fluorescence during calcium imaging. A thin layer of Chromium (~ 60nm) was sputtered on glass wafer before the first KMPR layer to block excitation light from reaching photoresist. The Cr was removed from the recording chamber region after the final RIE step with wet chromium etchant (MicroChem) to allow excitation light to reach only the *Hydra* in immobilization chamber. The Pt pads on the fabricated electrophysiology chip were connected to electrical leads with conductive epoxy. A microfluidic chip with similar immobilization chamber pattern (Fig. 2a) was aligned on top of the electrophysiology chip.

All microfluidic chips were reusable after cleaning. The microfluidic chips for electrophysiology and imaging were rinsed well with deionized water and oven dried at 80C for at least 40 minutes. The microfluidic chips used with chemicals were soaked in deionized water overnight (at least 10 hours) on a stir plate, sonicated in fresh deionized water for at least 10 minutes, heated to 160C for 1 hour, and finally oven dried at 80C for at least 40 minutes before reusing. We did not observe tentacle writhing-like behavior in the previously used devices (with GSH for feeding response) that were soaked in deionized water for at least 8 hours.

### *Hydra* Strains and Maintenance

The *Hydra vulgaris* AEP strains including the two transgenic *Hydra vulgaris* lines expressing either GCaMP6s in their neurons (GCAMP6s, neurons) and GFP in the neurons (GFP, neurons) were provided by Christophe Dupre in the laboratory of Rafael Yuste (Columbia University). All *Hydra* were cultured in *Hydra* Media using the protocol adapted from Steele laboratory (UC Irvine). *Hydra* were fed freshly hatched brine shrimp (artemia naupali) at least three times a week and the *Hydra* Media was replaced approximately 1-4 hours after feeding to remove excess food. The containers were thoroughly cleaned every four weeks to remove any film buildup. Individual *Hydra* were starved for at least 2 days prior to experiments with the exception of experiment with glutathione induced feeding behavior, where the animals were starved for at least 4 days.

### *Hydra* Loading and Unloading

*Hydra* was inserted into the microfluidic devices through a syringe cap attached to 1 mm tygon tubing that was inserted into the entry port. Using a glass pipette, *Hydra* was dropped into an open syringe cap then using the syringe connected to the port on the opposite end of the microfluidic immobilization chamber, negative pressure was used to pull the polyp into the immobilization chamber. During this process, the open syringe cap was connected to a syringe containing *Hydra* media to prevent inserting air into the device. In case when *Hydra* adhered to the tubing, alternating positive and negative pressures helped dislodge the *Hydra*. If *Hydra* still remained stuck, gentle localized tapping dislodged the *Hydra* to resume flow. This approach required working fairly quickly once the *Hydra* was dropped into the open syringe cap to prevent undesired adhesion to the plastic surfaces. Because of this stickiness, we had approximately 50% success rate for loading *Hydra* without causing significant damage. The second loading method reduced contact with plastic by pulling the *Hydra* few millimeters into the tubing with syringe then inserting the tubing into the inlet port of the microfluidic device. This approach increased success rate to 95%, though care had to be taken to not introduce any air into the microfluidic chamber. *Hydra* was loaded by applying positive pressure to the inlet syringe. The two opposing syringes were alternatively used to provide gentle pulses to position the *Hydra* at the recording site. At the end of the experimentation, *Hydra* could be removed from the microfluidic device either by disassembling the device or by gently pulsing the syringes to flow *Hydra* out of the large inlet port.

### *Hydra* Electrophysiology

Electrophysiology chip interfaced with PDMS was clamped with acrylic and the electrical leads were connected to the amplifier. All data was obtained with an Intan Technologies RHD2132 unipolar input amplifier (http://intantech.com) at a sampling rate of 10 KHz (for electrophysiology of *H. Vulgaris* AEP animals), low frequency cutoff and DSP filter of 0.1 Hz and high frequency cutoff of 7.5 KHz.

For electrophysiological experiments, *Hydra* starved for at least 48 hours was immobilized in the recording chamber and the recording began at least 5 minutes after *Hydra* had been immobilized. The animals were recorded from under ‘dark’ conditions with ambient light passed through red filter (Red filter #26, Roscolux). Six animals were recorded for one hour each (Supplementary Fig. 1) and three animals were recorded for 10 hours each (Fig. 2). The nano-SPEARs measured bursts of electrical activity when the animal contracted. These measurements resemble contraction bursts that are known to be associated with contraction. In cases when *Hydra* drifted away from the electrodes, we noticed decrease in signal amplitude. However, we could reestablish electrical contact by applying pressure from either the entry or the suction ports to reposition the animal.

Electrophysiology data had two obvious waveforms that correspond to behaviors: small spikes during tentacle contractions, large spikes during body contractions (Supplementary Fig. 1). The K-Means algorithm for clustering showed there were two optimal clusters. We manually selected spike amplitude threshold of 500 uV to derive the two distinct waveforms. In biphasic or triphasic waveforms, the largest negative or positive peak was used for the spike amplitude. Spike width was found by calculating the full width half max (FWHM) of the waveform. Successive large amplitude contraction pulses separated by 10 seconds or less were considered a part of the same contraction burst and the inter-pulse interval was the time between these pulses in a single contraction burst. Inter-burst interval was the time between contraction bursts.

### Simultaneous Imaging and Electrophysiology

Electrical measurements with nano-SPEARs were made while simultaneously performing brightfield or fluorescence imaging of either wildtype or transgenic animal, respectively. Brightfield imaging was performed (10 fps, 60 min, 0.13 N.A. 4x objective, and ‘dark’ lighting (see *Hydra* Electrophysiology methods) to record behaviors such as contractions occurring during measurement of electrical activity (Supplementary Movie 2). Transgenic *Hydra* expressing GCaMP6s in the neurons starved for at least 48 hours were used to measure the activity of the neurons (20 fps, 60 min, 0.45 N.A. 10x objective, and 20% light intensity with GFP filter) (Supplementary Movie 3). An inverted microscope equipped with GFP filter and Andor Zyla 4.2 were used for capturing all of the images. All electrical data was obtained with an Intan Technologies RHD2132 unipolar input amplifier (http://intantech.com) at a sampling rate of 1KHz, low frequency cutoff and DSP filter of 0.1 Hz and high frequency cutoff of 7.5 KHz.

### Movement Analysis for Simultaneous Electrophysiology and Imaging

Electrical activity from *Hydra* was recorded simultaneously with brightfield imaging. The images were binarized to extract *Hydra* regions using imbinarize and regionprop functionality in the Matlab Image Processing Toolbox. During body contractions, the fraction of the image corresponding to the *Hydra* body decreased, however, tentacle contractions were best quantified by measuring the area of the hypostomal region. Thus, the overall area occupied by the top half of the *Hydra* body (hypostome) was used to track *Hydra* movements because it could quantify both contraction bursts and tentacle movements (Fig. 2c). During body contractions, entire body including all tentacles contracted resulting in large decrease in hypostomal region area while during tentacles contraction, only few of the tentacles contracted resulting in smaller decrease in hypostomal region. Thus, increases or decreases in hypostomal area used to generate movement trace indicated when body and tentacle contractions occurred. To generate the movement maps (Fig. 2c, bottom right image in each box), we overlaid the binarized images of the *Hydra* and used the color map to represent the fraction of the time that the *Hydra* occupied each pixel. Thus, the light-colored areas of the map show the locations of body and tentacle contractions. Because *Hydra* remains contracted longer during closely occurring contraction bursts, dark-colored area of the map shows *Hydra* in its most contracted form (Fig. 2c left box). We used a 150 second time window to show body contractions more clearly because contraction bursts can last more than 30 seconds and sometimes do not elongate significantly between bursts. For correlation of tentacle contractions, we used a 30 second time window, which is comparable to the time scale of tentacle contractions.

### Correlation Analysis for Simultaneous Electrophysiology and Imaging

Electrical activity from transgenic *Hydra* (Neuronal, GCaMP6s) was recorded simultaneously with fluorescence imaging. From the calcium activity traces, we identified 30 sec intervals of either high amplitude activity or low amplitude activity to perform cross-correlation analysis. The high and low amplitude activity regions were manually identified with threshold of 20% of the highest peak in the calcium activity (Fig. 2c, d). The high amplitude activity region occurred during contraction bursts for neural activity imaging. The low amplitude activity region occurred during rhythmic potential like activity during neural imaging. For correlation maps (Fig 2 d), each frame was down sampled to 64 x 64 pixels and the vectors of fluorescence values for each of the down-sampled pixels across the 30 second interval were cross-correlated with electrical activity during the sample 30 sec interval using Matlab. The correlation value from each of the pixels was then used to determine the intensity of that pixel. The electrophysiological data were down-sampled and the fluorescence data were up-sampled to 100Hz for cross-correlation requiring vectors of equal length. Both the Intan amplifier and the Zyla were triggered with the same TTL signal. However, to account for any offset in the timing of the electrical and optical data, we measured the maximum of the cross-correlogram in 50 ms (approximately one duty cycle of the trigger signal) window rather than the cross-correlation at zero offset to generate the correlation maps.

### Fluorescence Imaging for Micro-Movement Analysis

Transgenic *Hydra* expressing GCaMP6s in the neurons was immobilized in an electrophysiology chip. *Hydra* movement was imaged with 488nm excitation laser and 0.45 N.A. 10x objective on Nikon Ti Eclipse Confocal microscope for 5 minutes at ~1 fps. Few cells with constant fluorescence were tracked for motion to calculate displacement and average movement of cells (65 um/minute) in the body column of *Hydra* immobilized in hour-glass chambers.

### Chemical Stimulation with Reduced Glutathione for Feeding Response

*Hydra* (*H. Vulgaris* AEP) were starved for at least 4 days prior to immobilization in the perfusion chambers for chemical stimulation. One side of the port connecting to the perfusion channels was used as the inlet port. Two syringes with stopcock valves containing either *Hydra* media or 9µM reduced glutathione (GSH) (Biosynth) were connected to a 2 to 1 manifold which was then connected to the perfusion input port. The inlet syringes were raised ~25 cm above the device to hydrostatically flow chemicals/buffer. Opposite side of the perfusion channels used as the outlet were connected to a syringe at the same height as the device. To calculate the flow rates into the observation chamber, we ignored all fluidic paths except narrow perfusion channels because the fluidic resistance in the narrow perfusion channels was significantly higher than in the tubing and thus had the largest contribution to flow rates. This calculation was in agreement with the approximately 0.02 mL/min change in syringe volume we observed during the experiments. After immobilization of the *Hydra*, different flow conditions were used while imaging behavioral changes for 30 minutes each. First, 30 minutes of baseline activity was fluorescently measured with no flow, followed by 30 minutes with flow of *Hydra* media to show minimal effects of slow perfusion on *Hydra* behavior. Next, perfusion input was switched to GSH for 30 minutes to cause the mouth to open and to inhibit body contractions. We began to measure response roughly 20 minutes after beginning the flow of GSH. Finally, the perfusion input was switched again to *Hydra* media for 30 minutes to terminate feeding response and recover normal contractile activity (Supplementary Movie 4). All brightfield imaging during chemical stimulation was performed using 0.13 N.A. 4x objective and Zyla4.2 at ~5 fps for 2 hours in 3 mm wide and ~160 µm tall observation chamber.

To obtain the *Hydra* length trace (Fig. 3c), we created binary images of the *Hydra* and measured the major axis length using Matlab image processing toolbox. During body contractions, body length decreases significantly as the *Hydra* contracts into a tight ball. The body length increases when the animal elongates. As a result, the traces of body length show large fluctuations indicating spontaneous body contractions for the first half of the experiment (60 minutes) and more constant body length once stimulated with GSH and mouth begins to open. The fluctuations in body length begin to return following wash with buffer. If the experiment is allowed to continue with wash buffer for another 30 minutes, the body contraction rate begins to approach the contraction rate prior to stimulation.

Transgenic *Hydra* (GCaMP6s, neurons) was starved for at least 4 days prior to immobilization and chemical stimulation. Fluorescence imaging during chemical stimulation was performed using 0.45 N.A. 10x objective, 12% epifluorescence light intensity and Zyla4.2 at ~15 fps for 1 hour in 3 mm wide and ~160 µm tall observation chamber. Because effective imaging time with fluorescence calcium indicator was close to thirty minutes before photobleaching occurred, the flow conditions were modified to reduce exposure to excitation light before chemical stimulation. First, the period of baseline activity measurement without flow was eliminated and period of *Hydra* media flow was reduced to five minutes. This was followed with GSH flow for thirty minutes, and finally recovery with *Hydra* media for thirty minutes (Supplementary Movie 5).

### Chemical Stimulation with Chloretone for Muscle Paralysis

Transgenic *Hydra* expressing GFP in the neurons was immobilized in the ~160 µm tall perfusion chambers for stimulation with 0.1% Chloretone (Acros Organics). The *Hydra* was imaged using 488nm excitation laser and 0.45 NA 10x objective and ~1fps on Nikon Ti Eclipse Confocal for tracking the neurons before and after being anesthetized. After perfusion of Chloretone, the animal movement was significantly decreased and the whole-brain anatomy was volumetrically imaged at high resolution with negligible motion artifacts (Fig. 1b).

### Behavior and Locomotion Tracking

*Hydra (H. vulgaris AEP)* raised in dark at 20C and fed three times a week under ambient light were used for behavioral and locomotive tracking. Time-lapse imaging was performed using USB digital microscopes. Specifically, we used either Dino-Lite Pro handled digital microscope (AM4114TL0M40, www.dinolite.us) or more affordable portable USB digital microscope/mini microscope endoscope (TOPMYS TM-M200, www.amazon.com). Images were captured at rate of 1 frame every 2 seconds on Windows built-in Camera application or open-source camera software iSpy (www.ispyconnect.com).

For behavioral tracking, evenly spaced 30 white LEDs were placed below a Roscolux diffuser for evenly illuminating behavioral microarena from the bottom (Fig. 4c left). *Hydra* was immobilized 2 days post feeding and imaged at 0.5 fps for 1 day at room temperature in roughly 440 µm tall microfluidic device with evenly spaced 2 × 1 mm pillars. The initial three hours of the recording in uniform lighting environment (n=1) were used for manually tracking the mouth, foot and body column center to quantify the *Hydra* position and posture (Supplementary Fig. 2). The Euclidean distances between the foot and the mouth were used to calculate the body length, L. The contraction bursts were identified as the minima in the body length values. Due to low frame rate, individual contraction burst pulses may not be sampled. To avoid counting multiple pulses within the same burst, minima with high prominence were identified for contraction bursts. The threshold for minima was determined from the average length of the *Hydra*. The angle of the vector from the foot to the mouth with respect to the positive x-axis was used to measure the body orientation, a, for exploration. The body curvature, β, was derived from the difference in angle between vector from center of the body to the mouth, α1, and the vector from foot to the center of the body, α2. Small β (β < π/3) indicated slight bend in the body while larger β (π/3 < β < π) indicated u-loop in the body posture. A translocation event was defined by displacement in foot position above a threshold with minimum time between such displacement events. The distance, d, and the direction, γ (with respect to positive x-axis), the *Hydra* moved was calculated from a vector from the previous position to new position of *Hydra* foot. Track of locomotion pattern was overlaid on the micrograph of *Hydra* in the microfluidic arena where the size of the nodes indicated the length of stationary periods and the length of the edges connecting the two nodes indicated displacement during translocation, d (number of pixels). The color map reflects the time from start of imaging where green is the start point (0 hours) and yellow is the end point (3 hours) (Fig. 4c). Raster plot of behavioral patterns was generated using above measurements (Fig 4c). Contraction burst duration was determined from the amount of time body length remained below a threshold. U-loop bend duration was determined by the amount of time body posture, b remained between π/3 and π. The translocation duration was determined by the amount of time it took for the displacement length, d, to return to baseline (defined by a lower threshold) before and after identified translocation step.

For behavioral tracking in environment with light gradient, white LED desk lamp light source pointed from the top towards one end of the microfluidic chip from above provided optical cue. For the purpose of identifying behavior influenced by light (n=1), the first 3 hours after immobilization were analyzed from one representative *Hydra* similarly to the *Hydra* in even illumination (see above) to generate individual behavioral plots (Supplementary Fig. 2), translocation map and raster plot of analyzed behavioral patterns (Fig. 4c right).

Phototactic locomotion was observed with *Hydra (n=5)* immobilized in the microfluidic chip for 10 – 72 hours. Some *Hydra* were successfully maintained in the microfluidic arena for up to 10 days but significantly decreased in size due to starvation. For phototactic locomotion, *Hydra* location was semi-automatically tracked by identifying the foot and its position (n=2) (Fig 4. d-f). First the images were binarized after background subtraction using Matlab image processing toolkit. *Hydra* foot was identified from binarized image using semi-automated algorithm that compared the extrema on each end of the major axis of *Hydra*. The end with lower variation in distance between extrema the body axis was identified as the foot. Tentacles were assumed to have greater spread in extrema of the binarized image. The algorithm required manual correction when the change in foot position was larger than a threshold in a single frame to correct for mislabeled foot position. The threshold was determined based on the size of the *Hydra*. A step function was fitted to the displacement over time to smooth it and reduce multiple small steps during translocation to a single step which was used to quantify the translocation events. Translocation maps (Fig. 4d) were generated using the displacement vectors similarly to the maps for 3 hours imaging (see above). Circular histogram of the translocation vectors was created by weighting direction of the vector by the length of the vector (Fig. 4e). Thus, large step in a specific direction meant higher preference for that direction compared to small step in another direction. Average translocation direction vector was calculated to indicate preference for translocation direction. The translocation direction preference was for the quadrants with brighter light illumination when analysis was performed on both the first three hours and first ten hours of immobilization. Circular histogram of body orientation, α, was created using unit vectors from *Hydra* foot to body centroid with respect to x-axis (Fig. 4f). Average body orientation vector was calculated for every image in the time-lapse to indicate preference for body orientation. The body orientation preference was not always in the same quadrants with brighter light illumination.

## ACKNOWLEDGEMENTS

We would like to thank Christophe Dupre and Dr. Rafael Yuste (Columbia University) for sharing the transgenic *Hydra* strains. This work is funded by the Defense Advanced Research Projects Agency Young Faculty Award D14AP00049 (J.T.R.), and the DARPA BioControl program (J.T.R). K.N.B. and D.L.G. are funded by training fellowships from the Keck Center of the Gulf Coast Consortia on the NSF Integrative Graduate Education and Research Traineeship (IGERT): Neuroengineering from Cells to Systems 1250104. D.L.G is funded by the NSF Graduate Research Fellowship Program 0940902. We also thank the Rice Shared Equipment Authority and the Nanofabrication Facility where the devices were fabricated.

## CONTRIBUTIONS

K.N.B. performed and analyzed experiments. K.N.B. developed the microfluidic platform for *Hydra*.

D.L.G., K.N.B. and D.G.V. developed the fabrication process for the nano-SPEAR electrophysiology chip.

B.W.A. provided hardware and software support. J.T.R. directed the research. K.N.B. and J.T.R. co-wrote the paper. All authors read and commented on the manuscript.

## Supplementary Figures

**Supplementary Figure 1:**
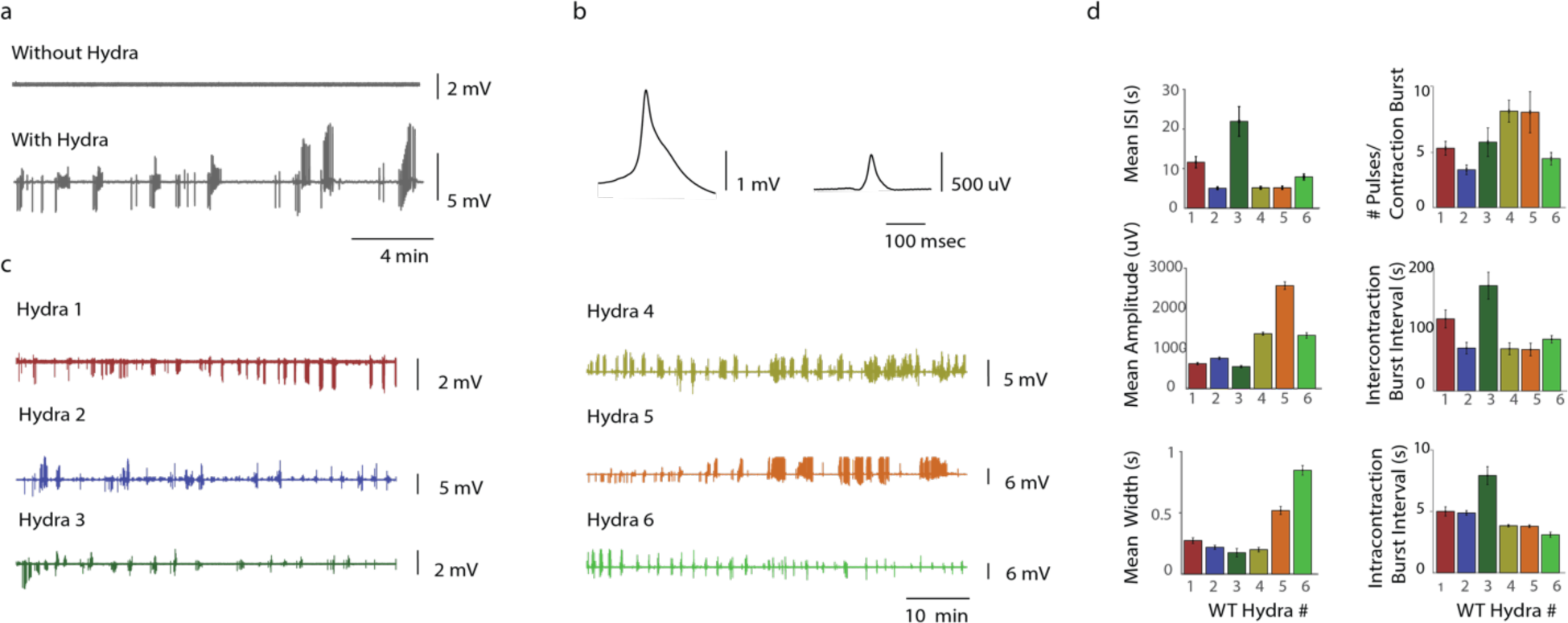
Analysis of electrophysiology of *Hydra*. **(a)** Shows 20 minutes electrical measurements from nano-SPEAR electrode with and without *Hydra* immobilized. **(b)** Peak-aligned average small and large amplitude waveforms determined by PCA clustering. **(c)** 1 hour long electrical recording from six individual *Hydra vulgaris* AEP. **(d)** Show contraction burst activity across six individual *Hydra* with analysis of large amplitude waveforms.

**Supplementary Figure 2:**
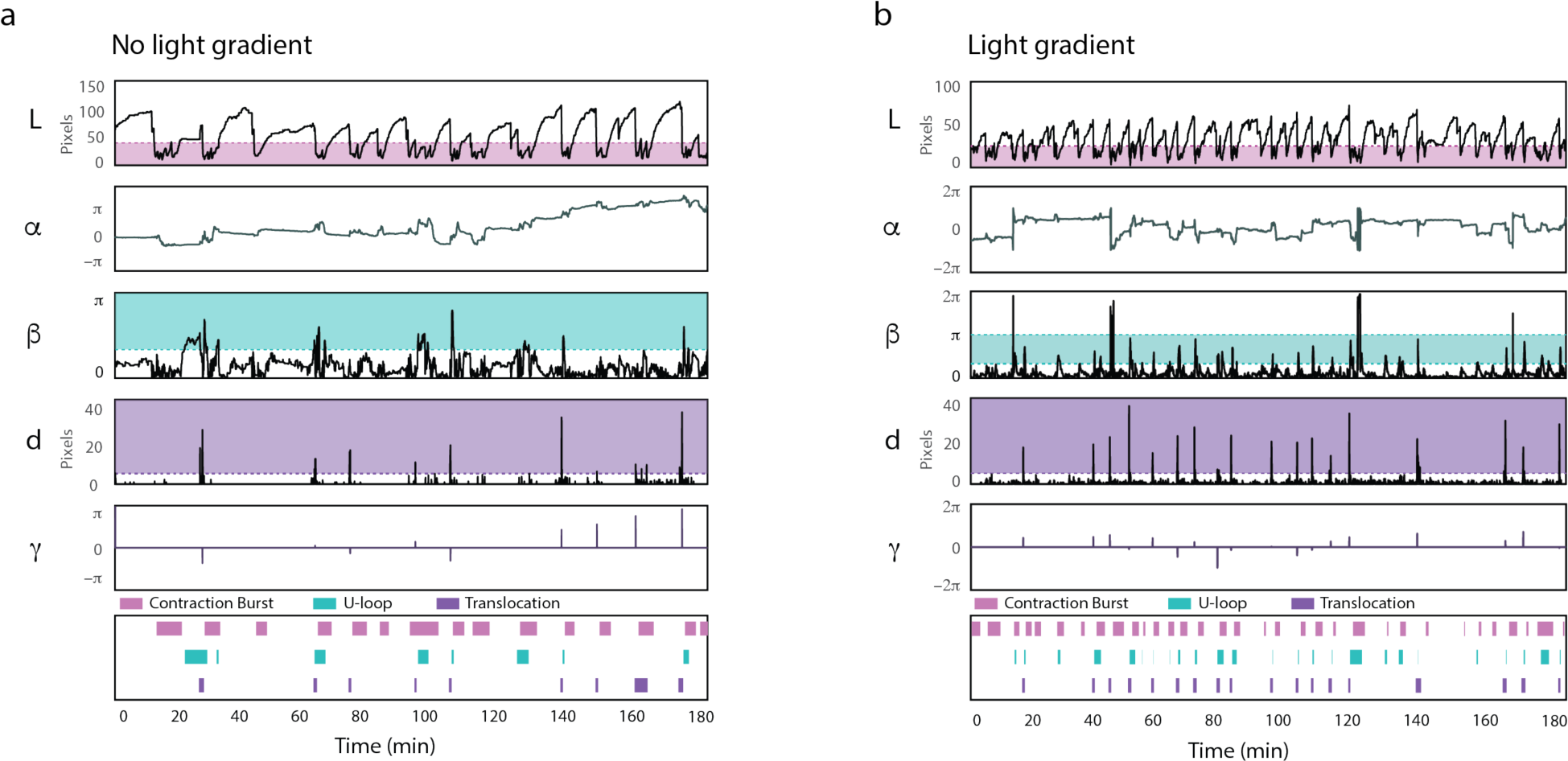
Behavioral analysis of *Hydra*. **(a)** Behaviors and locomotion pattern of *Hydra* under even illumination. **(b)** Behaviors and locomotion pattern *Hydra* under light gradient. (a,b) Plots show change in body length (L), body orientation (a), difference orientation of top half and bottom half of the body (b), displacement of foot (d) and direction of foot displacement (g). Shaded regions show threshold values used to generate raster plot. Contraction bursts are when body length falls within the shaded region. U-loop body bending is when b is within shaded region. Translocation event is when the displacement step size is within shaded region.

## Supplementary Movies

**Supplementary Movie 1: Microfluidic Immobilization of *Hydra***

Shows *Hydra* inserted through entry port and immobilized at the pinch point in hour-glass microfluidic device.

**Supplementary Movie 2: Simultaneous electrophysiology and brightfield imaging**

Shows simultaneous electrophysiology and brightfield imaging in *Hydra*. Trace at the bottom shows the electrical activity coinciding with movements.

**Supplementary Movie 3: Simultaneous electrophysiology and fluorescence imaging, neurons**

Shows simultaneous electrophysiology and calcium imaging of neurons in transgenic *Hydra*. Trace at the bottom shows the electrical activity coinciding with observed calcium activity.

**Supplementary Movie 4: Brightfield imaging of mouth opening during chemical stimulation**

Shows chemical stimulation of *Hydra* in wheel-and-spoke device. Labels at the top indicate flow conditions: i) no flow, ii) media flow, iii) GSH flow, and iv) media flow. Tentacles start to contract half way into GSH flow and mouth cavity later forms. *Hydra* loses rigidity after mouth opening and is easier to push aside with minimal flow conditions. Mouth begins closing and contractions slowly return half way into media flow following stimulation.

**Supplementary Movie 5: Calcium imaging of mouth opening during chemical stimulation**

Shows fluorescence imagine of calcium indicator in neurons during chemical stimulation of transgenic *Hydra* (GCaMP6s, neurons) in wheel-and-spoke device. Labels at the top indicate flow conditions: i) media flow, ii) GSH flow, and iii) media flow.

**Supplementary Movie 6: Time-lapse imaging of *Hydra* behavior in 2D**

Time-lapse movie shows *Hydra* moving in evenly illuminated microfluidic arena. Simplified movement track is overlaid to show detected translocation events.

